# Prediction and Characterization of Disorder-Order Transition Regions in Proteins by Deep Learning

**DOI:** 10.1101/2021.06.11.448022

**Authors:** Ziang Yan, Satoshi Omori, Kazunori D Yamada, Hafumi Nishi, Kengo Kinoshita

## Abstract

The biological functions of proteins are traditionally thought to depend on well-defined three-dimensional structures, but many experimental studies have shown that disordered regions lacking fixed three-dimensional structures also have crucial biological roles. In some of these regions, disorder–order transitions are also involved in various biological processes, such as protein-protein interaction and ligand binding. Therefore, it is crucial to study disordered regions and structural transitions for further understanding of protein functions and folding. Owing to the costs and time requirements of experimental identification of natively disordered or transitional regions, the development of effective computational methods is a key research goal. In this study, we used overall residue dependencies and deep representation learning for prediction and reused the obtained disordered regions for the prediction of disorder–order transitions. Two similar and related prediction tasks were combined. Firstly, we developed a novel deep learning method, Res-BiLstm, for residue-wise disordered region prediction. Our method outperformed other predictors with respect to almost all criteria, as evaluated using an independent test set. For disorder-order transition prediction, we proposed a transfer learning method, Res-BiLstm-NN, with an acceptable but unbalanced performance, yielding reasonable results. To grasp underlining biophysical principles of disorder-order transitions, we performed qualitative analyses on the obtained results and discovered that most transitions have strong disordered or ordered preferences, and more transitions are consistent with the ordered state than the disordered state, different from conventional wisdom. To the best of our knowledge, this is the first sizable-scale study of transition prediction.

**Availability:** https://github.com/Yanzziang/Transition_Disorder_Prediction

## Introduction

Intrinsically disordered proteins or regions do not form stable structures under physiological conditions and yet have crucial roles in biological processes, including the recognition of proteins, nucleic acids, and other types of molecules, post-translational modifications, and alternative splicing (Dyson and Wright, 2005; Nishi, et al., 2013; Tompa, 2005; Uversky and Dunker, 2010; Uversky, et al., 2008; Wright and Dyson, 1999). Some of these disordered proteins or regions undergo a transition from the disordered to ordered state (or vice versa) upon interactions with other molecules. In particular, segments that transition from disorder to order via protein–protein interactions are recognized as molecular recognition features (MoRFs) and have been explored extensively (Kotta-Loizou, et al., 2013; Mishra, et al., 2018; Mohan, et al., 2006; van der Lee, et al., 2014; Yan, et al., 2016).

In response to the increasing importance of intrinsically disordered regions and disorder–order transitions, numerous disordered region predictors have been developed in the last two decades (Liu, et al., 2019). These predictors are roughly divided into physicochemical property-based approaches (such as IUPred (Dosztanyi, et al., 2005) and FoldIndex (Prilusky, et al., 2005)) and machine learning-based approaches (such as DisEMBL (Linding, et al., 2003), SPINE-D (Zhang, et al., 2012), SPOT-Disorder (Hanson, et al., 2017),and SPOT-Disorder-single (Hanson, et al., 2018)). These software and servers, particularly with machine-learning based approaches, have achieved fairly good accuracy, which allows us to explore potentially disordered regions without waiting for experimental verification. Similarly, predictors of functional sites in disordered regions have also been developed and the majority of these employ protein-interacting MoRFs as training datasets (Disfani, et al., 2012; Hanson, et al., 2019).

Even though it is widely accepted that disorder-order transitions can occur not only by protein interactions but also by small ligand binding, allosteric effects, and other environmental changes, not many prediction studies have been performed on non-MoRF transitions. Furthermore, disordered protein regions have recently been recognized as druggable regions (Ruan, et al., 2019), implying that drugs should be designed to cause disorder–order transitions to work effectively. To fully understand or control the functions and dynamic behaviors of protein disordered regions, a more general understanding of disorder–order transitions and transition regions is essential.

In this study, we developed a disorder–order predictor, Res-BiLstm, by employing state-of-the-art machine learning techniques, such as the bidirectional long short-term memory (LSTM) with residual connections. LSTM, a variant of recurrent neural networks (RNNs), can deal with contextual information for input data and thus are widely utilized for biological sequence analyses (Jiang, et al., 2018; Liu and Gong, 2019; Yamada and Kinoshita, 2018). Our predictor exhibited equivalent or superior performance to those of existing methods. We then expanded our method to predict transition sites recorded in the Protein Data Bank (PDB). This method, Res-BiLstm-NN, inspired by the effectiveness of transfer learning in MoRFs, enabled the application of disorder information to the prediction of transitions and exhibited acceptable but unbalanced performance. An exploratory analysis using our predictors revealed that transition sites have unique features compared to those of non-transition (order or disorder) regions, different from the conventional view. To the best of our knowledge, this is the first attempt to capture the general characteristics of transition sites without assumptions regarding molecular interactions.

## Methods

### Datasets

#### Disordered protein dataset

A dataset consisting of 4229 disordered proteins from SPINE-D (Zhang, et al., 2012) was used. A total of 3000 proteins were selected for training, 400 proteins were selected for validation in the learning process, and the remaining 829 proteins were used as an independent test set. The disordered protein dataset was extremely unbalanced; disordered regions only occupied approximately 10% of all disordered proteins. Table S1 enumerates the unbalanced disordered and ordered residues in training, validation, and test datasets.

#### Transition site dataset

To build the transition site dataset, all human enzymes stored in the Universal Protein Resource (UniProt) (UniProt Consortium, 2018) were retrieved. All identified protein structures of human enzymes (Du, et al., 2014) were extracted from PDB (Berman, et al., 2002), and the ss_dis.txt file (Berman, et al., 2002) was used to obtain notations about the order/disorder states. Then, summary statistics about structural information at the residue level was obtained and residues were classified as transition, non-transitional ordered, or disordered sites. First, the residues of all extracted X-ray structures from PDB were aligned to the corresponding UniProt amino acid sequences with reference to the SIFTS file (Dana, et al., 2019; Velankar, et al., 2013). Sequence segments whose structural information were not recorded in any structure files were removed. Order/disorder labels indicated that the residues were always observed as ordered/disordered among protein structures for the protein. The transition label indicated that the residues were identified as both ordered and disordered at least once among available protein structures. Proteins without transition labels were excluded, resulting in 2995 proteins in the final dataset. Figure S1 summarizes the process of building the dataset. The human protein dataset was split into 2200 proteins for training, 300 proteins for validation, and 495 proteins for the independent test set. The dataset was unbalanced. Residues labeled as transition, non-transitional order, and disorder occupied 75,793, 828,617, and 59,560 residues in the human protein dataset respectively (Table S2), where the residues located within three residues of the N-terminus or C-terminus were excluded from statistical analyses.

### Model features

There are 20 kinds of amino acids constituting variable-length protein sequences. To obtain the amino acid characters, similar to SPINE-D, evolutionary information, physicochemical properties, and predicted structural information were used as features for each amino acid within a protein, instead of sequence-based information. The 20 evolutionary features were extracted from a position-specific scoring matrix (PSSM), which was generated by three iterations of a PSI-BLAST search (Altschul, et al., 1997) against the non-redundant protein sequence dataset UniRef50 (Suzek, et al., 2015).

Seven physicochemical properties defined by Meiler et al. (Meiler, et al., 2001), including van der Waals volume, hydrophobicity, isoelectric point, and so on, were also used to characterize amino acids. In addition, 17 predicted structural features, such as probabilities of secondary structures, torsional angles, contact number, and so on, were generated using SPIDER2 (Heffernan, et al., 2015). These parameters were used to generate a 44-length feature vector for each amino acid. As a distinction, in three-state classification, these features were particularly scaled from 0 to 1 by the min-max normalization method. Figure S2 shows the feature matrices for different protein sequences.

### Neural network architectures

#### Architecture of disordered region prediction

Unlike standard feedforward neural networks, like multilayer perceptron (MLP), RNNs can deal with variable-length input sequences by memory functions for previous input information. In particular, LSTM (Gers, et al., 2000; Hochreiter and Schmidhuber, 1997) and bidirectional LSTM (Graves and Schmidhuber, 2005) have excellent learning performance. The bidirectional LSTM improves performance by reading input sequences in both forward and backward directions simultaneously. Owing to its mechanism, the bidirectional LSTM is essentially a deep network architecture and could be difficult to be optimized due to the gradient varnishing or exploding problem. To resolve this issue, residual learning, which adopts shortcut connections to perform identity mapping (He, et al., 2016), was introduced. In this study, the bidirectional LSTM layers and residual connections were combined to obtain disordered region predictions, in an approach termed stacked <u>Res</u>idual <u>Bi</u>directional <u>LSTM</u> network, or Res-BiLstm. In brief, Res-BiLstm consists of five bidirectional LSTM layers with concatenated output vectors from forward and backward LSTM blocks. Residual connections were added from one layer to the next. In the network, the sigmoid unit is adopted at the last discriminated layer for structural binary classification of each amino acid. The layout of the Res-BiLstm network is shown in Fig. S3. In the architecture, 100 neural units are set inside the recurrent hidden layers of LSTM blocks. An input protein sequence is 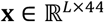, where L and 44 indicate the length of the protein sequence and number of input features for an amino acid, respectively. Input biological features for position *t* within the protein sequence, 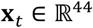, are represented layer-by-layer and transformed to a fixed vector 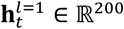 -by performing the following forward transformations on **x**_*t*_:

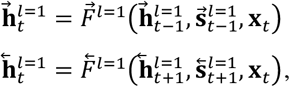

where arrow, **h, s**, *l* = *a*, and *F* are forward or backward direction, the output vector, internal memory of the LSTM cell, *a*-th layer, and gating functions of LSTM blocks-in the forward and backward direction. The concatenated output vector 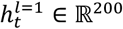 is calculated by the following equation:

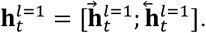

Thus, the output vector from fifth LSTM layer, 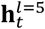 is calculated as follows:

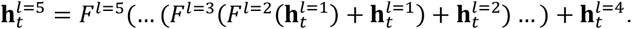

Finally, the representation vector **x**_*t*_ and 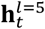 are passed into the sigmoid function for structural binary classification. The Res-BiLstm model is trained by a weighted binary cross entropy function to assign each amino acid to one of two classes, disordered or ordered, as follows:

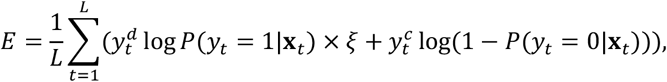

where 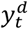 and 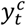 represent the corresponding class label of disordered and order regions and the class weighting ξ is added to account for unbalanced classes. In our problem, the weight, ξ is set to 8.853 for the disordered class based on the ratio of the two classes among all amino acids of training set.

#### Prediction of the architecture of transitions

Apart from natively disordered and ordered regions, proteins can also contain regions undergoing a spontaneous disorder-order transition. MoRFs are disordered regions undergoing a unidirectional disorder-to-order transition while binding to partners. Previous computational prediction methods for MoRFs show relatively good performance using biological features extracted from protein sequences (Sharma, et al., 2018). A transfer learning method based on predictions of disordered regions showed improved MoRF identification (Hanson, et al., 2019). These results suggest that disorder-order transition regions can be predicted based on sequence information, and knowledge of disorder is transferable for the prediction of transition regions. In this study, a transfer learning method was implemented to bridge two similar tasks, residue-wise disordered region prediction and transition prediction. Our aim was to improve transition identification learning based on the transition dataset we generated by transferring learned knowledge for a sequence-to-structure mapping based on the disordered protein dataset.

Similar to the architecture of previous MoRF prediction, we extracted latent representations from the pre-trained disordered region predictor Res-BiLstm as input for the transition prediction task. As theoretically demonstrated in the previous subsection, the Res-BiLstm network with a constant error carousel in each LSTM block can effectively overcome gradient vanishing and exploding, which adequately maintains the actual dependencies among amino acids within a sequence. Deep stacking with residual connections between each LSTM layer enables more powerful characterization through layer-by-layer disorder encoding. The last disorder representation containing a high level of disorder–order structure information is reused for further transition encoding. The architecture of the transition prediction is shown in Fig. S4. Our method, Res-BiLstm-NN, involves the last representation layer of the pre-trained predictor Res-BiLstm and an MLP including three fully connected hidden layers with 100 nodes and dropout (ratio of 0.2) in each layer. As in Res-BiLstm, the sigmoid function is used at the last discriminant layer for the binary classification.

In the model, 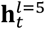 is used to execute transition encoding, which can be described by the following equations:

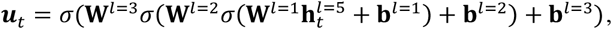

where, σ represents ReLU, 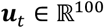 is the encoded vector from the last fully connected layer, and **W** and **b** are parameters to be learned.

Finally, in the encoded vector **x**_*t*_, **u**_*t*_ is passed to a sigmoid decision unit for binary transition classification. Similar to Res-BiLstm, the error is calculated by a weighted binary cross entropy by the following equation:

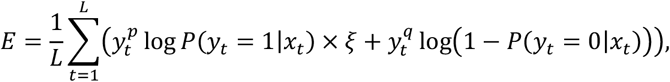

where 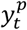 and 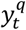 represent the corresponding class labels for structural transitions and non-transitions. The class weigh ξ is set to 11.3 for unbalanced classes.

#### Optimizer and regularization

The two models Res-BiLstm and Res-BiLstm-NN were trained by the Adam optimizer with a learning rate of 0.001. Early stopping was executed when the validation error stops decreasing in the learning process to reduce the risk of overfitting.

#### Computational environment

The training processes were implemented using the NVIDIA V100 Graphics Processing Unit for rapid training on the NIG supercomputer at the ROIS National Institute of Genetics. As a framework for neural network implementation, Keras version 2.2.4 and Tensorflow version 1.13.1 were utilized as back-end software.

## Results

### Disordered region prediction

#### Performance evaluation

The performance of Res-BiLstm was evaluated using the test dataset. For comparison, we also evaluated the performance of IUPred (long) and IUPred (short), DisEMBL, SPINE-D, and SPOT-Disorder. As a criterion, we utilized the area under the receiver operating characteristic (ROC) curve (AUC). As shown in Fig. 1, Res-BiLstm achieved the highest AUC for the independent test dataset, indicating the effectiveness and superiority of stacked residual LSTM networks. AUC values were similar for Res-BiLstm and SPOT-Disorder (differing by 0.0003), implying that stacked residual representation learning does not provide a significant benefit compared with a shallow-layers BiLstm method. SPINE-D was comparable to our method, and this can likely be explained by the common dataset used for network training. In addition to the AUC value, other metrics were employed for the balanced and comprehensive measurement of model performance. We used sensitivity and specificity as metrics of the ability to correctly identify disordered and ordered residues, respectively. Owing to unbalanced classes in the disordered protein dataset, traditional accuracy metrics are not directly applicable. Instead, we used a balanced accuracy measurement (ACC) by the calculating mean of the sensitivity and specificity values. We applied various metrics to compare the performance of several disordered region predictors, as shown in Table 1. Res-BiLstm and SPOT-Disorder outperformed other predictors with respect to almost all metrics, including SPOT-Disorder-Single, which utilized a common dataset for network training. This superior performance is likely due to the additional effective evolutionary, structural, and physicochemical features compared with pure sequence information. The performance of SPINE-D, utilizing nearly identical features to those of Res-BiLstm, was relatively weaker than that of our methods, implying that a sliding window approach restricts the discriminant analysis of sequences to structures. In addition to AUC, Res-BiLstm exhibited a much higher sensitivity than that of SPOT-Disorder, thereby yielding the best ACC, indicating that this method exhibits the most balanced performance for correctly identifying both disordered and ordered regions. MCC and F1 values for our method were second only to SPOT-Disorder; it is likely that the higher specificity of SPOT-Disorder plays an important role in unbalanced datasets containing large samples of ordered regions and small samples of disordered regions.

**Fig. 1.**
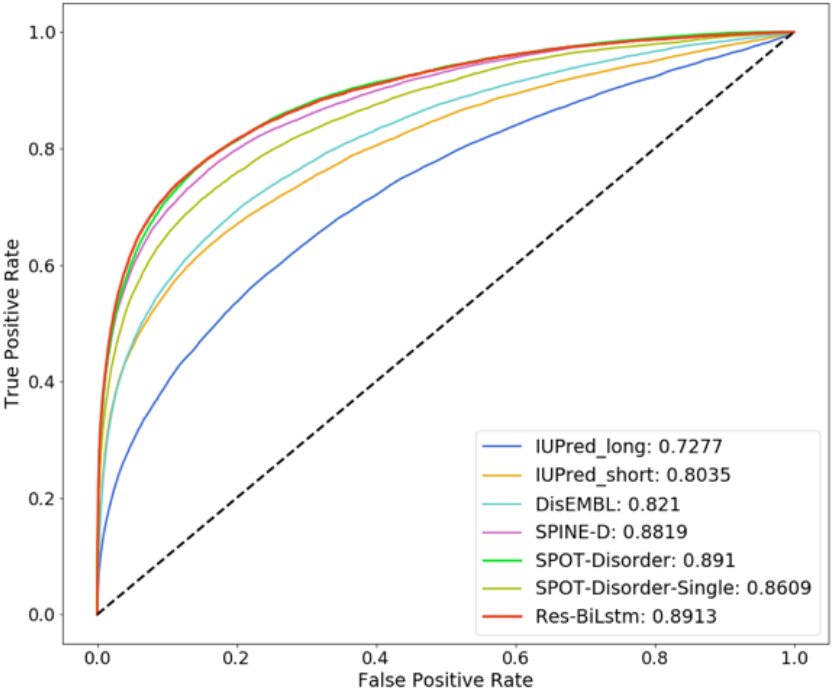
Comparison of sensitivity, specificity, and AUC values for several disordered region predictors

**Table 1.** Comparison of major metrics among several predictors.

|  | AUC | ACC | Se | Sp | MCC | F1 |
| --- | --- | --- | --- | --- | --- | --- |
| IUPred (long) | 0.728 | 0.619 | 0.283 | 0.955 | 0.267 | 0.318 |
| IUPred (short) | 0.804 | 0.706 | 0.466 | 0.946 | 0.403 | 0.454 |
| DisEMBL | 0.821 | 0.706 | 0.463 | 0.950 | 0.411 | 0.461 |
| SPINE-D | 0.882 | 0.801 | **0.748** | 0.854 | 0.420 | 0.447 |
| SPOT-Disorder | **0.891** | 0.736 | 0.493 | 0.978 | **0.541** | **0.567** |
| SPOT-Disorder-Single | 0.861 | 0.676 | 0.365 | **0.987** | 0.486 | 0.485 |
| Res-BiLstm | **0.891** | **0.808** | 0.702 | 0.913 | 0.493 | 0.529 |

The maximum value in each column is shown in bold. Here, ACC, Se, Sp, MCC and F1 stands for accuracy, sensitivity, specificity, Mathew's correlation coefficient and F1 score.

#### Example of disorder prediction outputs

To determine actual output probabilities for an entire protein sequence, we selected an experimentally annotated protein from the UniProt database (ID: P03176), thymidine kinase. We obtained the prediction probability curves for thymidine kinase from several disordered region predictors, as shown in the Fig. S5.

Res-BiLstm obtained a more stable output than those of predictors based on energy estimation methods. Moreover, the output probabilities obtained using Res-BiLstm included more extreme values (extremely high for the actual disorder class or low for the actual order class) than those of predictors that learn discriminant patterns using other computational methods. Based on these results, our method can more effectively identify both disordered and ordered regions.

### Transition prediction

#### Performance evaluation

The performance of the transition region predictor Res-BiLstm-NN was evaluated on the test dataset. The confusion matrix for the binary classification problem and evaluation metrics are shown in Fig. S6 and Table 2, respectively. As shown in Table S3, our method identified disordered and ordered regions with high accuracy. However, more than half of the transitions were incorrectly classified as structurally non-transitional states, and the ratio of correctly predicted transitions and incorrectly predicted non-transitional regions made it difficult to correctly identify transitions using Res-BiLstm-NN. As presented in Table 2, based on AUC and ACC values, our method exhibited acceptable performance in general. The low MCC and F1 values indicated an unbalanced performance for detecting transition regions. These results reflect the difficulty in predicting transition regions.

**Table 2.** The performance of Res-BiLstm-NN.

|  | AUC | ACC | Se | Sp | MCC | F1 |
| --- | --- | --- | --- | --- | --- | --- |
| Res-BiLstm-NN | 0.741 | 0.641 | 0.432 | 0.849 | 0.194 | 0.259 |

#### Example of predicted transition sites

We selected DNA polymerase kappa (UniProt ID: Q9UBT6) to analyze the actual output probabilities of Res-BiLstm-NN,.

As presented in Fig. S7, almost all of the transition sites were correctly predicted. However, a number of structurally unchangeable regions were also incorrectly classified, suggesting that some may undergo a potential transformation in specific environmental conditions. Interestingly, the output probability curve exhibits relatively drastic fluctuations even among residues in close proximity, implying that transitions are likely to be determined by or influenced by specific environmental conditions than by their original inherent structures.

The three-dimensional structure of the DNA polymerase kappa protein is presented in Fig. 2. It was found that one of the correctly predicted regions (shown in blue) is closely located to thymidine triphosphate, suggesting that our predictor successfully identified the disorder-order transition region which may associated with ligand binding and then protein function.

**Fig. 2.**
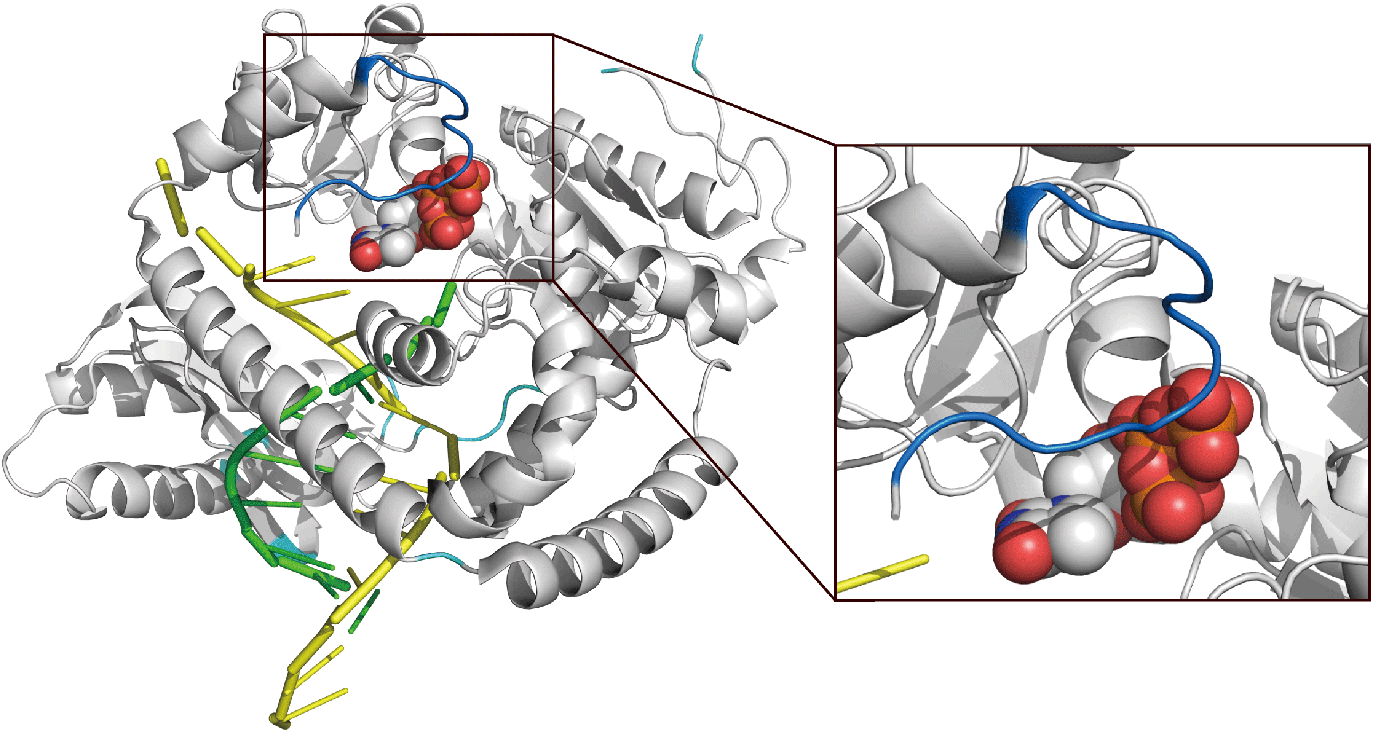
Three dimensional structure of DNA polymerase kappa (UniProt ID: Q9UBT6, PDB ID: 2OH2 chain B). DNA strands are shown in yellow and green. The ligand (thymidine triphosphate) is shown as sphere. The correctly predicted transition regions are shown in cyan and blue. It is suggested that the disorder-order transition of the blue region might associate with ligand binding.

## Discussion

We developed an accurate predictor of disordered protein regions, Res-BiLstm, and applied it to the first large-scale analysis of structural transitions based on a transfer learning method, Res-BiLstm-NN, which uses the internal representation of protein disorder from Res-BiLstm. The AUC values for an independent test dataset indicated an acceptable overall performance but values for sensitivity, MCC, and F1 highlight the difficulty in identifying the transition class. This can potentially be explained by the strong disorder–order preference, potential unknown transitions, and inherently inseparable sequence information distribution.

We applied Res-BiLstm, which produces disordered or ordered predictions for each residue, to the transition site test dataset. This process allowed us to explore the similarity between transitions and natively disorder/order sites. We also performed disordered region prediction using the architecture of Res-BiLstm trained on the order-disorder-transition dataset without transition sites. The results are shown in Fig. 3. The probability distribution predicted by the established predictor is presented in the left-hand panel. A relatively small fraction of transitions was in an intermediate state, and the majority showed a strong disorder or order preference. The output distribution from direct learning (right-hand panel) shows that a large proportion of transitions were classified as ordered regions. Compared to an intermediate state between disorder and order, most transition sites had strong disorder or order preference. Thus, the inseparable similarity due to the strong disorder-order preference can negatively impact model performance. More structural transition sites are likely to be similar to ordered sites than to disordered sites, whereas substantial evidence shows that transition regions prefer the disordered states for entropic advantage when they are in ligand free states, and the number of known disorder-to-order transitions associated with ligand binding far exceeds the opposite cases in common datasets, such as DisProt (Uversky, 2002).

**Fig. 3.**
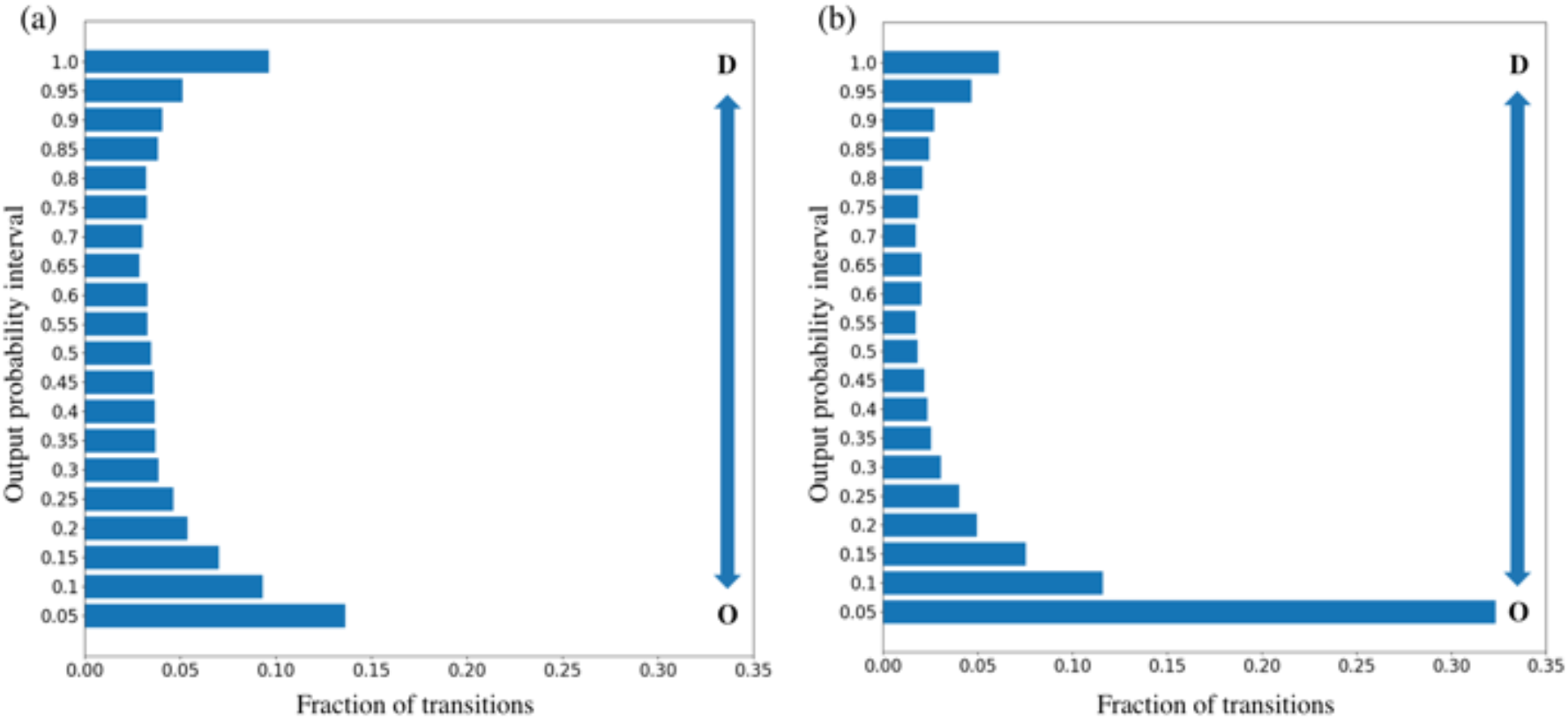
Predicted output distributions of transitions. Disordered region prediction by (a) Res-BiLstm and (b) same architecture in direct learning.

To further explore transition preferences, we performed a three-state (transition, non-transitional order, and non-transitional disorder) classification based on the transition dataset. We employed a similar architecture to Res-BiLstm, but used the softmax function in the last discriminant layer instead of the sigmoid function. As presented in Fig. S8, the low sensitivity of transitions indicates that transition site prediction from sequence information alone is difficult. In other words, evolutionary, structural, and physicochemical information extracted solely from protein sequences might not be sufficient to identify whether a region can undergo a structural transformation or not. Furthermore, transition sites tend to be misclassified to non-transitional ordered sites more frequently than to non-transitional disordered sites. This indicates that transition sites are similar to ordered sites than to disordered sites, to a certain degree. Table S4 shows various metrics for individual binary-class evaluations. Disorder-Others performed better than Order-Others with respect to most metrics, such as AUC, ACC, and MCC, supporting the similarity between transition sites and ordered sites, different from the conventional view.

## Conclusion

We developed two accurate protein disordered region predictors, Res-BiLstm and Res-BiLstm-NN. Our predictors showed decent performance for predicting ordered/disordered and transition regions. Exploratory analyses using our predictors revealed that transition sites have unique features compared to non-transition (ordered or disordered) regions. To the best of our knowledge, this is the first attempt to capture the general characteristics of transition sites without assumptions regarding molecular interactions.

## Acknowledgements

Computations were partially performed on the NIG supercomputer at ROIS National Institute of Genetics and on the ToMMo supercomputer at Tohoku Medical Megabank Organization.

## Funding

This work was supported in part by Platform Project for Supporting in Drug Discovery and Life Science Research (Basis for Supporting Innovative Drug Discovery and Life Science Research (BINDS)) from AMED under Grant Number JP19am0101067.

## Conflict of Interest

none declared.

## Notes

https://github.com/Yanzziang/Transition_Disorder_Prediction

### Competing Interest Statement

The authors have declared no competing interest.

