## Supplemental file for "Prediction and Characterization of Disorder-Order Transition Regions in Proteins by Deep Learning"

### Supplementary Figures and Tables for Prediction and Characterization of Disorder-Order Transition Regions in Proteins by Deep Learning

Ziang Yan[1], Satoshi Omori[1], Kazunori D Yamada[1], Hafumi Nishi[1] and Kengo Kinoshita[1,2,3] *

[1]Graduate School of Information Sciences, Tohoku University, Sendai, 980-8579, Japan

[2]Tohoku Medical Megabank Organization, Tohoku University, Sendai, 980-8573 Japan.

[3]Institute of Development, Aging, and Cancer, Tohoku University, Sendai, 980-8575 Japan.

*To whom correspondence should be addressed.

**Supplementary Table S1.** Unbalanced number of disordered and ordered residues

| | Proteins | Ordered residues | Disordered residues |
|---|---|---|---|
| Training set | 3000 | 656634 | 74170 |
| Validation set | 400 | 86463 | 11721 |
| Testing set | 829 | 190285 | 17361 |

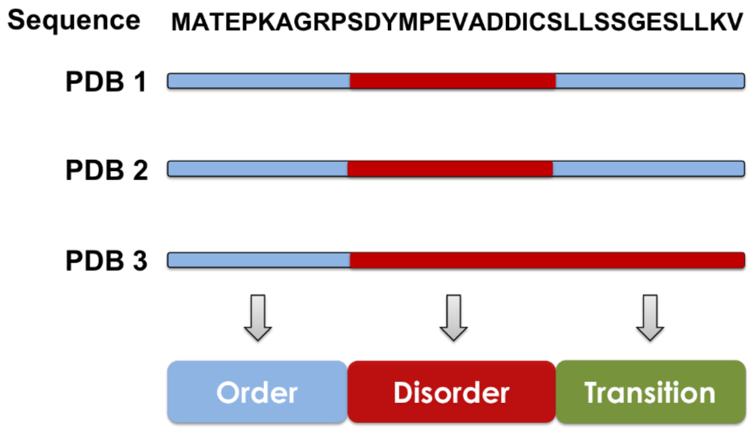


**Supplementary Fig. S1.** Classification scheme of transition sites at the residue level. Sequence: Amino acid residues of a human enzyme in Universal Protein Resource. PDB n: PDB structures of the enzyme. Order/disorder notation of each residue is shown in cyan (order) and red (disorder). Classification of transition is based on the statistics of these notations. The transition label indicated that the residues were identified as both ordered and disordered at least once.

**Supplementary Table S2.** Unbalanced residues labeled as transition, non-transitional order and disorder.

| | Proteins | Ordered | Disordered | Transition |
|---|---|---|---|---|
| Training set | 2200 | 624571 | 43539 | 58727 |
| Validation set | 300 | 77168 | 5622 | 6142 |
| Testing set | 495 | 126878 | 10399 | 10924 |

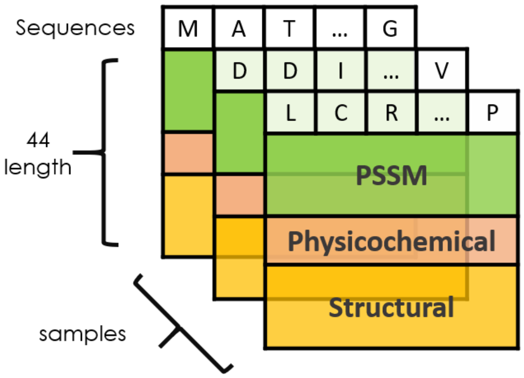


**Supplementary Fig. S2.** Feature matrices of protein sequences. The feature vectors consist of 20 evolutional features from PSSM, 7 physicochemical features, and 17 predicted structural features.

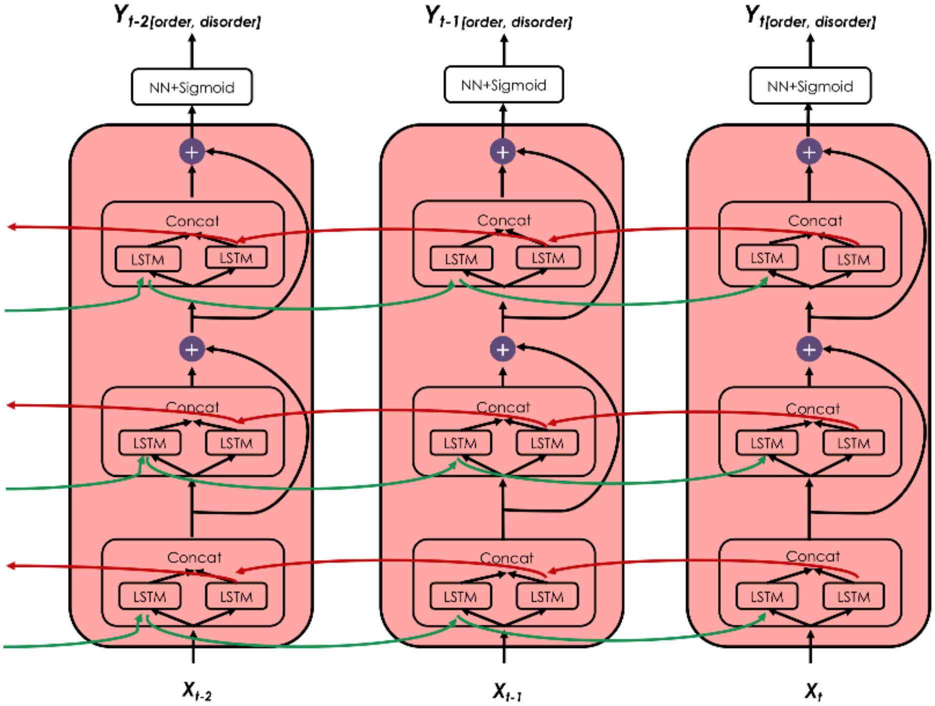


**Supplementary Fig. S3.** Layout of Res-BiLstm network. $\mathbf{x}_t$ is feature vector of a residue at position $t$. $\mathbf{y}_t$ is the output probability for a residue at position $t$. Probability less than 0.5 or more than 0.5 is classified to order or disorder respectively.

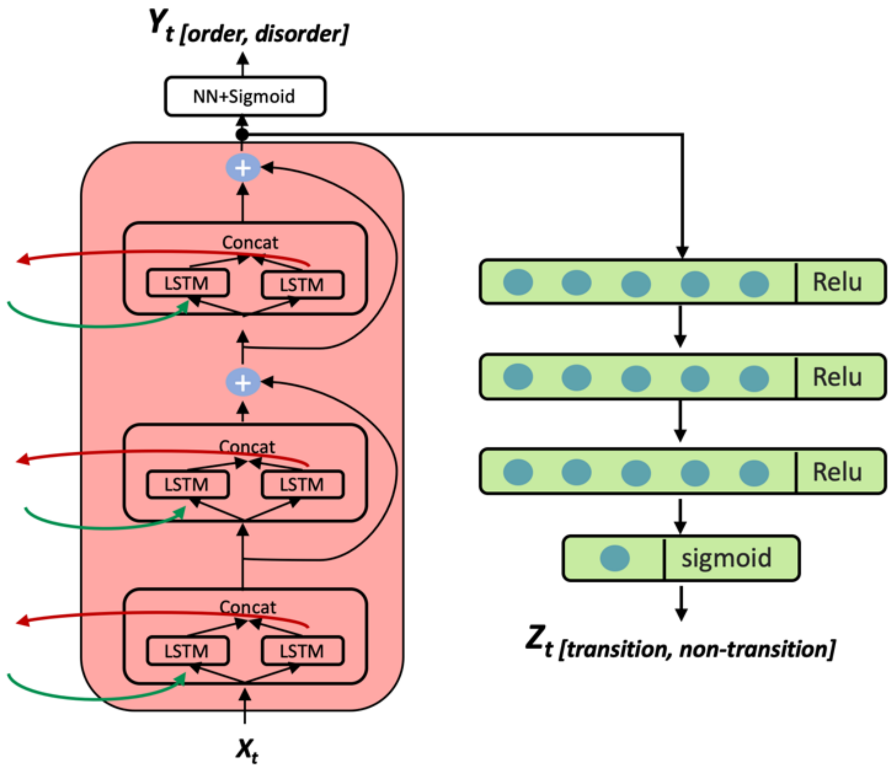


**Supplementary Fig. S4.** Layout of Res-BiLstm-NN network. $\mathbf{x}_t$ is feature vector of a residue at position $t$. $\mathbf{y}_t$ is output probability of Res-BiLstm and is shown here standing for a disorder characterization transfer from Res-BiLstm. $\mathbf{z}_t$ is the output probability for a residue at position $t$ from Res-BiLstm-NN. Probability less than 0.5 or more than 0.5 is classified as non-transition site or transition site.

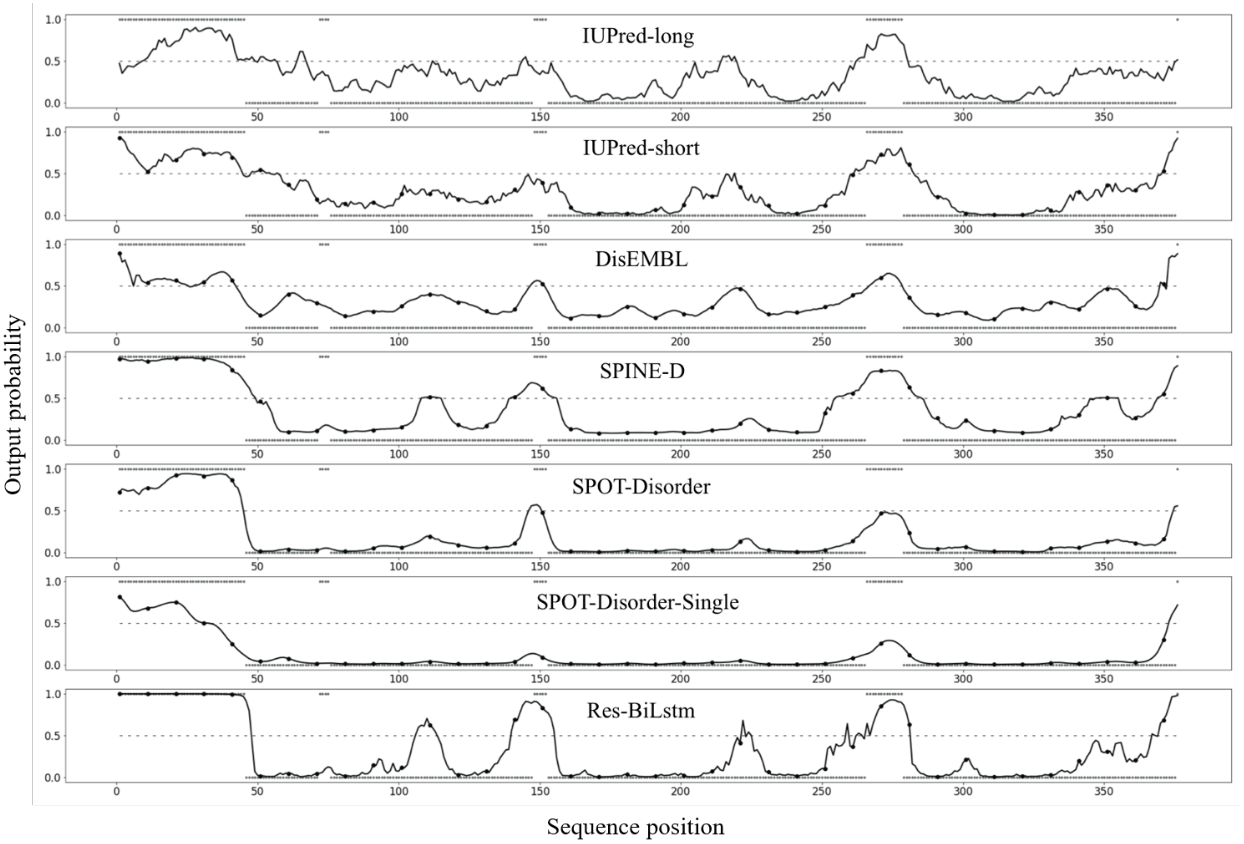


**Supplementary Fig. S5.** Comparison of output probabilities of several disordered region predictions for Thymidine kinase (UniProt ID: P03176). Each point in the curve represents the predicted probability for a particular protein sequence position. An output probability of 0.5 is the threshold for discriminating disordered (>0.5) or ordered (<0.5) classes. The dots on the top and bottom of each plot represent the actual disordered labels and actual ordered labels respectively.

| Ture label \ Predicted label | Transitions | D&O |
|---|---|---|
| Transitions | 4721<br>(Sensitivity: 0.43) | 6203 |
| D&O | 20797 | 116480<br>(Specificity: 0.85) |

**Supplementary Fig. S6.** Confusion matrix of residue-wise prediction by Res-BiLstm-NN.

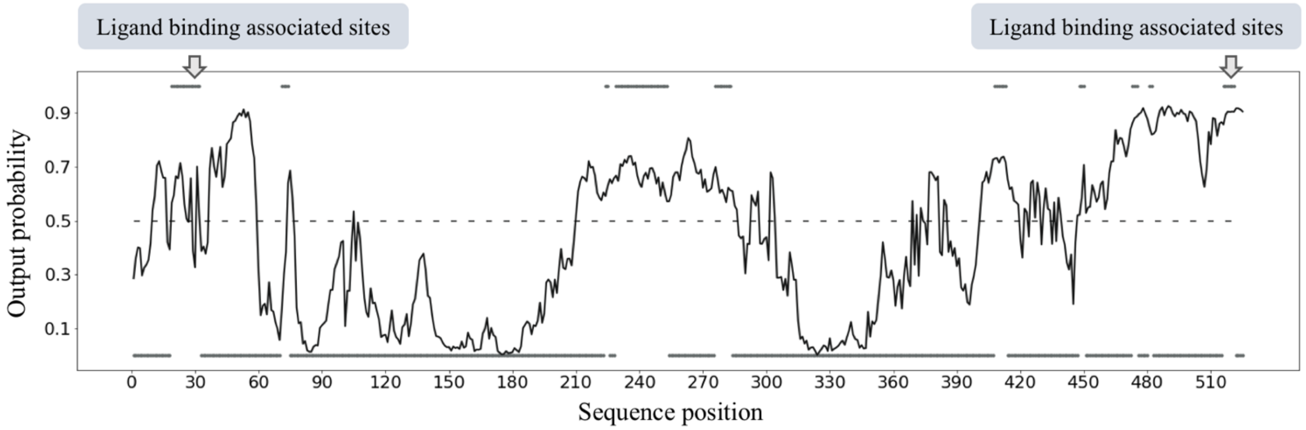


**Supplementary Fig. S7.** Output probabilities of transition regions for protein DNA polymerase kappa (UniProt ID: Q9UBT6) predicted by Res-BiLstm-NN. Each point in the curve represents the predicted probability for a particular protein sequence position. Dotted line stands for a threshold of 0.5 for discriminating transition (>0.5) or non-transition (<0.5) classes. The dots on the top and the bottom of the plot represent the actual transition labels and actual non-transition labels respectively. Original sequence has 870 residues, but tail-end 345 residues which contains a large amount of unobserved structure are not presented here.

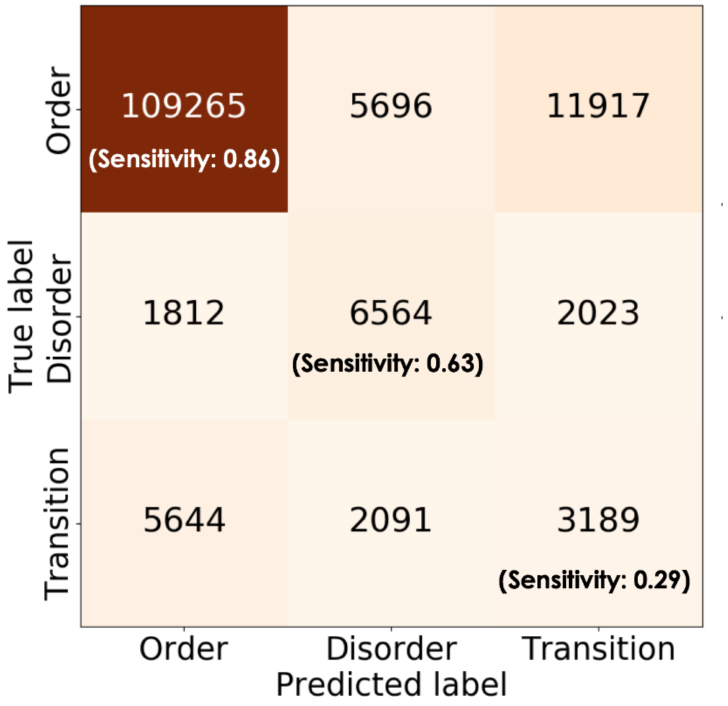


**Supplementary Fig. S8.** Confusion matrix of three-stage classification.

**Supplementary Table S4.** Major metrics of individual binary-class evaluations.

| Evaluation | AUC | ACC | Se | Sp | MCC | F1 |
|---|---|---|---|---|---|---|
| Order-Others | 0.839 | 0.756 | **0.861** | 0.650 | 0.439 | **0.897** |
| Disorder-Others | **0.908** | **0.787** | 0.631 | **0.943** | **0.496** | 0.530 |
| Transitions-Others | 0.711 | 0.595 | 0.292 | 0.898 | 0.156 | 0.227 |
